# Perinatal ischemic stroke impairs motor and cognitive development, muscle structure, and gut microbiota composition, with functional improvement following fecal microbiota transplantation in a mouse model

**DOI:** 10.64898/2026.06.23.733929

**Authors:** Gianmarco Cuboni, Catalina Campuzano, Livia Vignozzi, Rosa Liotta, David Pinzauti, Giampiero Vitale, Marika Tonellato, Roberto di Gesù, Manuele Biazzo, Michela Rigoni, Manuela Allegra, Gabriele Deidda

**Affiliations:** Fondazione di Ricerca Pediatrica (IRP), Padua, Italy; Department of Biomedical Science, University of Padua, 35121 Padua, Italy; Padova Neuroscience Center, University of Padua, 35121 Padua, Italy; Neuroscience Institute, National Research Council (IN-CNR), Padua, Italy; IRCCS ISMETT, Palermo, Italy; UPMC ITALY, Palermo, Italy; The BioArte Limited, Life Science Park, SGN300, San Gwann; Malta; Musculoskeletal Tissue Engineering (MsTE) lab, Ri.MED Foundation, Palermo, Italy; Faculty of Medicine & Dentistry, Queen Mary University of London, Malta Campus, VCT 2520 Victoria, Malta

## Abstract

Perinatal ischemic stroke is an early developmental brain injury caused by obstruction of cerebral blood vessels and is a leading cause of cerebral palsy and cognitive disability in survivors. However, progress in understanding its impact on the brain and other organ systems, as well as in developing effective therapies, remains limited, in part due to the scarcity of relevant preclinical models. Here, we induced ischemic stroke via middle cerebral artery occlusion in perinatal mice and investigated its effects within and beyond the brain across development into adulthood. We found that perinatal stroke disrupted fine motor development and impaired memory. In addition, it induced structural alterations in skeletal muscle and significant changes in gut microbiota composition. Notably, gut-targeted intervention using fecal microbiota transplantation improved fine motor function. Our findings demonstrate, for the first time, the multisystem developmental impact of perinatal stroke, extending beyond the brain, and identify gut microbiota modulation as a promising and potentially safe therapeutic strategy to improve motor outcomes after stroke.

## INTRODUCTION

Perinatal ischemic stroke is a brain injury occurring early in development, caused by blockage of cerebral blood vessels, and represents the main cause of cerebral palsy and cognitive impairments in survivors (1,2). In humans, perinatal stroke occurs between 28 weeks of gestation and 28 days after birth and accounts for approximately 25% of all strokes occurring during development, with an estimated incidence of 1 in 1600 live births (3); however, the true incidence may be underreported due to diagnostic challenges in many regions (4). Although several studies reported long-term neurological consequences of perinatal stroke affecting both motor and cognitive functions (5,6), most investigations focused on single post-injury time points. As a result, a comprehensive understanding of how perinatal stroke influences functional outcomes across development and into adulthood remains lacking. At the molecular level, signaling pathways involved in neurodevelopment and post-ischemic repair may contribute to both central and peripheral consequences of stroke. One such pathway is mediated by CXCR4 (C-X-C chemokine receptor type 4), a rhodopsin-like G protein-coupled receptor that binds the chemokine CXCL12. CXCR4-CXCL12 signaling plays critical roles in neurogenesis, immune cell trafficking, and post-ischemic tissue remodeling in adult stroke models (7,8). However, its role and expression following ischemic injury in the developing brain and its potential contribution to long-term extracerebral outcomes remain poorly defined. Motor dysfunction is among the most prevalent and persistent sequelae of stroke. In humans, approximately 80-85% of stroke-related deficits involve the motor system (9). While fine motor impairments following adult stroke are well documented (10), perinatal stroke likewise results in profound deficits that disrupt motor development, academic performance, and quality of life (11,12). Affected children frequently exhibit muscle weakness, with or without spasticity, often presenting as hemiplegia (12). In preclinical settings, rodent models enable reliable assessment of these motor deficits using standardized behavioral paradigms, providing a platform for mechanistic and therapeutic studies (13). Beyond central nervous system injury, ischemic stroke in adulthood exerts widespread systemic effects, including alterations in skeletal muscle structure and function and profound changes in gut microbiota composition and diversity (14,15). In contrast, the effects of perinatal stroke on skeletal muscle and other peripheral components of the motor system, as well as on the microbiota, despite the high prevalence of lifelong impairment in this population (4). Moreover, the progress in the development of new effective therapies is limited by the scarcity of preclinical models and a general lack of awareness and research focus on this critical issue (16,17). Currently, therapeutic options remain limited (18,19). Recent work explored novel intervention strategies, including approaches targeting the gut-brain axis based on the alterations in gut microbiota composition and stroke severity and functional recovery. Microbiota-based interventions, such as fecal microbiota transplantation (FMT), have been shown to reduce infarct volume and improve neurological outcomes in adult rodents (20,21). More broadly, FMT demonstrated protective effects across a range of neurological and systemic pathologies in both preclinical and clinical contexts (19,22). These considerations raise several key questions. First, how perinatal stroke affects long-term motor and cognitive function remains incompletely understood. In parallel, it is unknown whether perinatal ischemic injury results in persistent alterations in skeletal muscle structure and other motor-related peripheral tissues, or whether early-life stroke is sufficient to induce long-lasting changes in gut microbiota composition and diversity. At the mechanistic level, the involvement of molecular pathways known to regulate recovery after adult stroke, particularly CXCR4 signaling, has yet to be elucidated in the context of perinatal stroke. Finally, from a translational perspective, it remains unclear whether therapeutic strategies targeting the gut-brain axis are effective after perinatal stroke. Specifically, whether microbiota-based interventions such as FMT can modulate central and peripheral outcomes, improve motor or cognitive function, or reduce lesion volume following perinatal ischemic injury has not yet been investigated. Here, we induced stroke in mice at perinatal age by means of the distal middle cerebral artery occlusion (dMCAO) model (23) and we studied the impact inside and outside the brain. We found that perinatal stroke reduces neuronal density and induces brain atrophy, accompanied by changes in CXCR4 expression across the acute and chronic phase. Perinatal stroke disrupted fine motor function throughout development into adulthood and impaired memory performance. In addition, it caused structural alterations in skeletal muscle and significant shifts in gut microbiota composition. Notably, early gut-targeted intervention via fecal microbiota transplantation (FMT) after stroke improved fine motor function in adulthood. Collectively, these findings provide a comprehensive view of the long-term impact of perinatal stroke and identify gut microbiota modulation as a promising therapeutic strategy for improving fine motor outcomes after stroke.

## RESULTS

### dMCAO in P14 mice models perinatal stroke

Since the perinatal period in mice spans from birth to postnatal day 15 (P15), and stroke induction in this period results in a low survival rate (24), we induced ischemic stroke by dMCAO at P14 (**Fig. 1a**) to ensure high survival rates while remaining within the perinatal developmental window (25,26). dMCAO successfully produced a *core* lesion in the first and secondary somatosensory cortices (**Fig. 1a**, Supplementary **Fig. 9a**), providing a closer approximation to human stroke lesions (27,28). First, we quantified the extent of brain damage at 2 and 79 days after stroke induction (D2 and D79). At D2, dMCAO induced a *core* lesion of 0.559 ± 0.001 mm^3^ that corresponded to 7.35% of the ipsilateral hemisphere (**Fig. 1b**), with no differences between stroke and sham animals in terms of brain (**Fig. 1d**) or hippocampal (**Fig. 1f**) atrophy. In adulthood, at D79, we found that lesion volume stood at 5.792 ± 0.001 mm³ that correspond to 6.788% of the ipsilateral hemisphere (**Fig. 1c**), and stroke animals showed brain atrophy (-7.56 ± 0.01 %) in comparison to sham-operated mice (**Fig. 1e**), with negligible hippocampal atrophy (**Fig. 1g**). These data show that dMCAO at P14 successfully recapitulates the brain lesion pattern of perinatal ischemic stroke.

**Figure 1.**
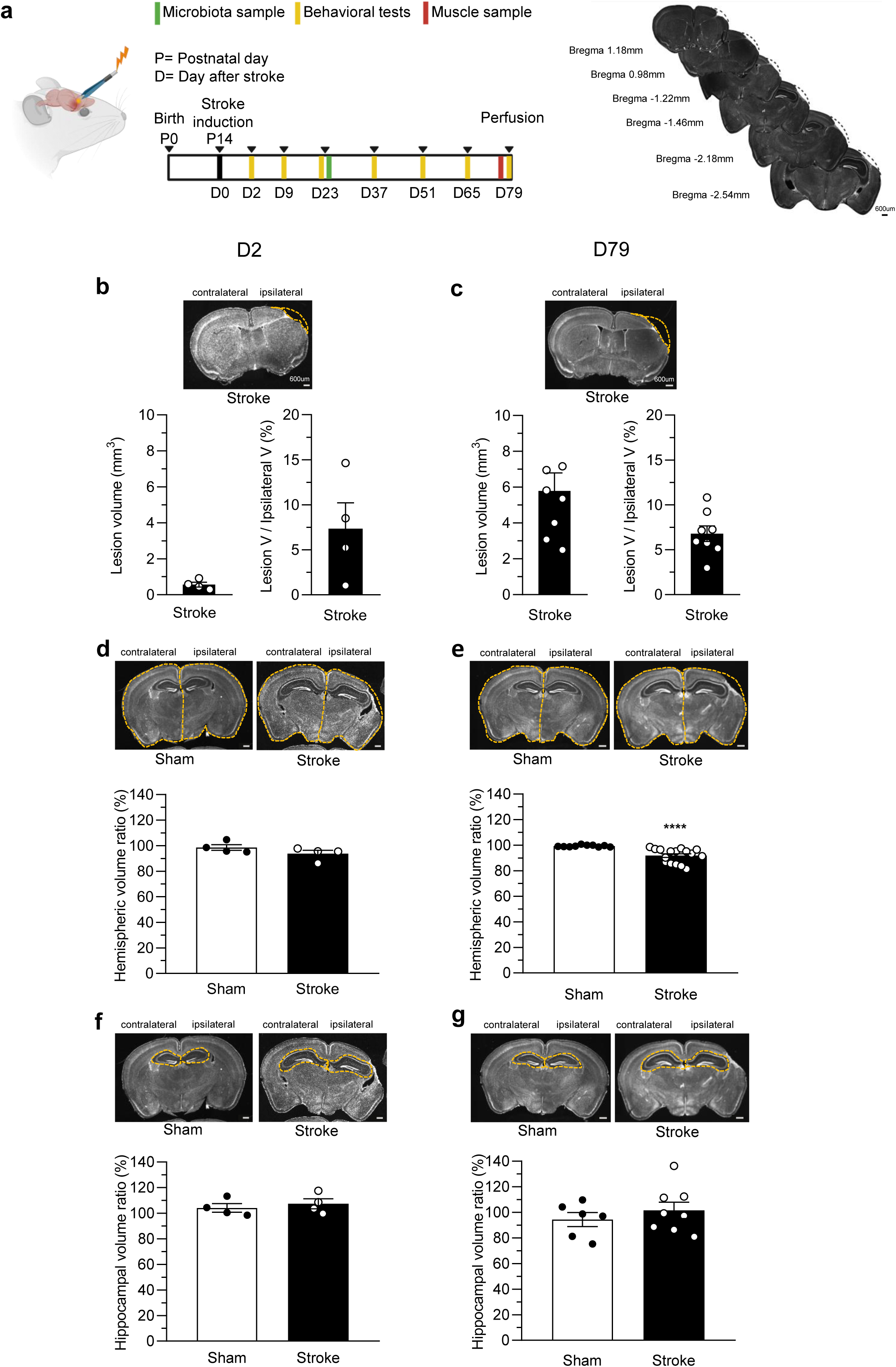
dMCAO in P14 mice models perinatal stroke. (**a**) Left, schematic cartoon of the experimental protocol. Right, representative coronal sections of the mouse brain 79 days after stroke induction with corresponding coordinates. (**b,c**) Top, representative coronal section of a mouse brain 2 days (**b**) and 79 days (**c**) after perinatal stroke. Bottom, the lesion volume quantification in stroke animals, 0.559 ± 0.001 mm³, with the percentage of the lesion (7.35%) compared to the ipsilateral hemisphere at D2 (**b**) and at d79 after stroke induction, 5.792 ± 0.001 mm³, 6.788% (**c**). (**d**,**e**) Top, representative coronal sections of the ipsilateral and contralateral hemispheres 2 days (**d**) and 79 days after perinatal stroke (**e**). Bottom, brain atrophy quantification showing no difference between sham (white) and stroke (black) animals at D2 (Student’s t-test; P = 0.1989), while a reduction after 79 days between sham (white) and stroke (black) animals (P = 0.0003). (**f**,**g**) Top, representative coronal sections of the ipsilateral and contralateral hippocampus 2 days (**f**) and 79 days (**g**) after perinatal stroke. Bottom, quantification of the hippocampal atrophy seeing no differences between in sham (white) and stroke (black;animals at D2 (Student’s t-test; P = 0.5542) (**f**) and in sham (white) and stroke (black) at d79 (Student’s t-test; P = 0.4299) (**g**). All the D2 images have a 200 μm scale bar. All the D79 images have a 600 μm scale bar. Scale bars of 200 μm (D2) and 600 μm (D79) are labeled only in the first image, while the following images display unlabeled scale bars. All histograms show mean ± s.e.m. (See Methods for details on animal numbers) Statistical significance *P<0.05, **P<0.01, ***P<0.001.

### Perinatal Stroke Alters Neuronal Survival and CXCR4 expression

To assess neuronal loss and CXCR4 expression following perinatal ischemic stroke, we performed immunohistochemical analyses at D2 and at D79, focusing on the penumbral region surrounding the core lesion. Neuronal loss was quantified by measuring neuronal density using the anti-NeuN antibody. Quantification of NeuN-positive cell density revealed a significant reduction in stroke mice compared to sham controls at D2 (**Fig. 2a**), whereas a more pronounced decrease was observed at D79, indicating progressive neuronal loss over time (**Fig. 2d**). No significant differences were detected between the perilesional and mediolateral regions within the lesioned hemisphere of stroke mice (data not shown). CXCR4-positive cell density was increased in stroke animals relative to sham controls at D2 (**Fig. 2b**), but significantly reduced at D79 (**Fig. 2e**). Consistently, CXCR4 signal intensity in neuronal processes (identified by neurofilaments labeling, NF) is higher in the stroke group at both time points (**Fig. 2c,f**), more markedly at D2. Collectively, these findings indicate that perinatal ischemic stroke induces progressive neuronal loss and dynamic changes in CXCR4 expression in the peri-infart region. Neuronal loss, early CXCR4 upregulation in neuronal processes, and its progressive decline at chronic stages recapitulate findings reported in adult ischemic stroke models (29).

**Figure 2.**
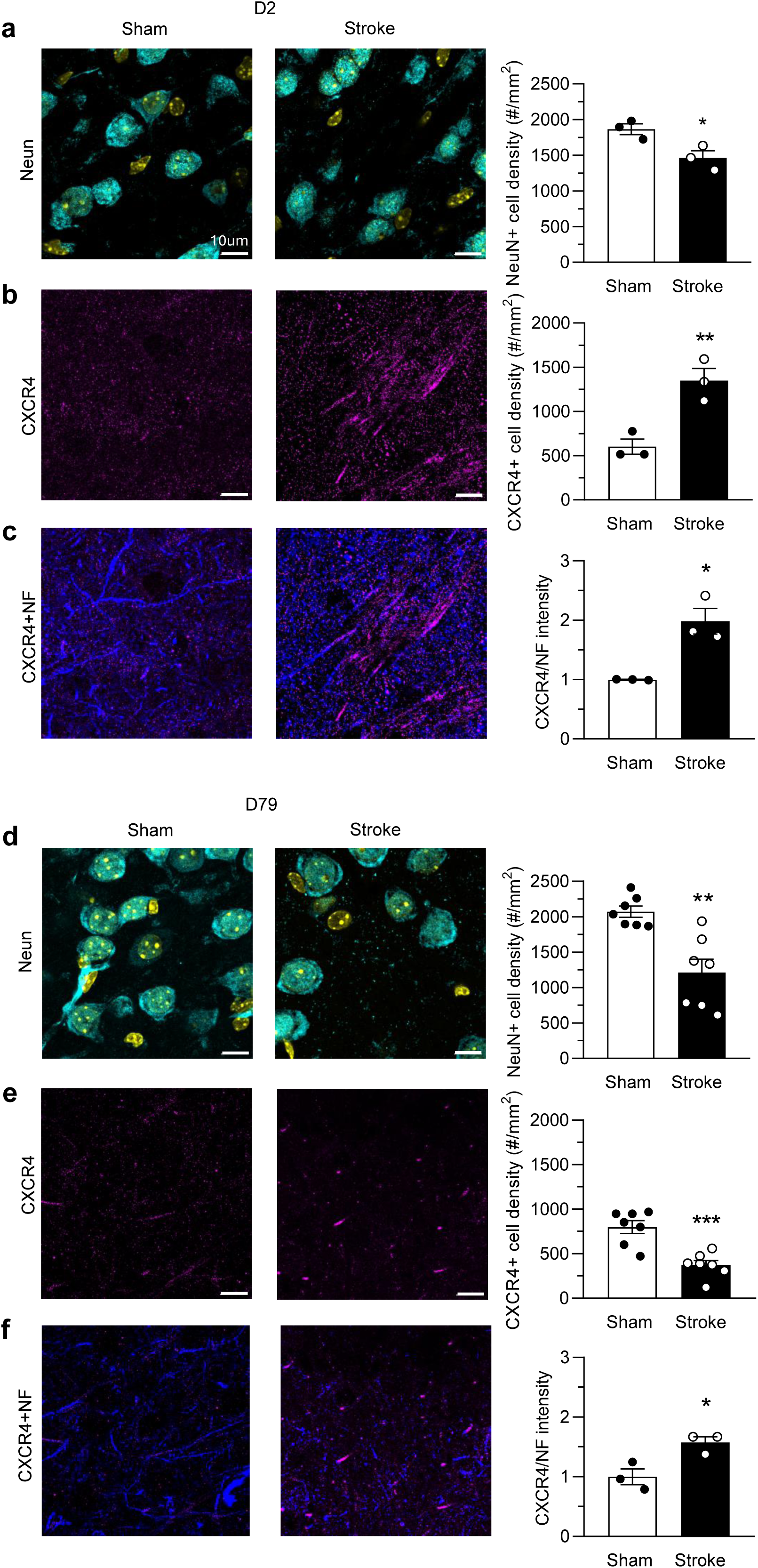
Perinatal stroke alters neuronal survival and CXCR4 expression. (**a**) Left, representative confocal images of visual cortex sections from sham and stroke animals 2 days after perinatal stroke (D2), immunostained with anti-NeuN (cyan), and DAPI (yellow). Right, quantification of NeuN-positive cell density at D2. A significant reduction was observed in the stroke group vs sham animals (Student’s t-test, P = 0.0325). (**b**) Left, representative confocal images of visual cortex sections from sham and stroke animals at D2, immunostained for CXCR4 (magenta). Right, quantification of CXCR4-positive cell density at D2, which was significantly increased in the stroke group (Student’s t-test, P < 0.001; P = 0.0099) (**c**) Left, representative confocal images of visual cortex sections from sham and stroke animals at D2, immunostained with anti-CXCR4 (magenta) and anti-Neurofilament (NF, blue) antibodies. Right, quantification of CXCR4 intensity in NF-positive processes, which is significantly increased in the stroke group (data normalized on sham) (Student’s t-test, P < 0.01; P = 0.0105). (**d**) Left, representative confocal images of visual cortex sections from sham and stroke animals 79 days after perinatal stroke (D79), immunostained with anti-NeuN (cyan) and DAPI (yellow). Right, quantification of NeuN-positive cell density at D79, which is reduced in the stroke group (Student’s t-test, P < 0.01 P = 0.0013). (**e**) Left, representative confocal images of visual cortex sections from sham and stroke animals at D79, immunostained with anti-CXCR4 (magenta). Right, quantification of CXCR4-positive cell density at D79, which was significantly decreased in the stroke group (Student’s t-test, P < 0.0001; P = 0.0005). (**f**) Left, representative confocal images of visual cortex sections from sham and stroke animals at D79, immunostained with anti-CXCR4 (magenta) and anti-NF (blue). Right, quantification of CXCR4 signal intensity in NF-positive processes at D79, whose intensity is higher in the stroke group vs sham, though to a less extent than at D2 (data normalized on sham) (Student’s t-test, P < 0.01; P = 0.0253). All the images have a 10 μm scale bar. The scale bar is labeled only in the first image, while the following images display unlabeled scale bars. All histograms depict mean ± s.e.m., and circles represent individual animals. (See Methods for details on animal numbers). Statistical significance *P<0.05, **P<0.01, ***P<0.001.

### Perinatal ischemic stroke impacts fine motor behavior

In this study, the dMCAO-induced stroke left the primary motor cortex as a perilesional *penumbra* (30) which plasticity process may drive the motor recovery after adult (31,32) and perinatal stroke (33), thereby enabling to testing of the motor functional outcome. Specifically, we tested the motor outcome longitudinally after dMCAO or sham surgery on different days (D; 2, 9, 23, 37, 51, 65, and 79) after stroke induction (**Fig. 3a**). To assess motor coordination and balance, we performed the rotarod test (see Methods; **Fig. 3b**, top panel)(13). Across all the developmental ages tested (D2-D79), stroke mice performed similarly to sham mice, suggesting that perinatal stroke did not impair overall motor coordination and balance (**Fig. 3b**, bottom panel). Next, we evaluated the muscle strength of the upper limbs by means of the grip strength test (see Methods; **Fig. 3c**, top panel) (34). We found no significant differences between stroke and sham mice (**Fig. 3c**), indicating that perinatal stroke did not affect the upper limb strength. To assess fine motor development and performance, we took advantage of the gridwalk test (13), already widely used to unmask fine motor impairments after adult stroke (35) (**Fig. 3d**, upper panel) (13). We quantified foot faults in the left forelimb, as it was contralateral to the lesion in the right hemisphere (36). In particular, we analyzed the asymmetry index (see Methods). Interestingly, stroke mice showed an increased asymmetry index in all the experimental days *versus* sham mice (**Fig. 3d**, bottom panel), indicating a higher number of foot faults, mirroring a deficit in fine motor movements induced by perinatal stroke. Since previous studies showed differences in the degree of recovery outcomes within a stroke sample (37,38), we differentiated the stroke group between good and poor spontaneous recoverers, namely, mice that recovered and mice that did not recover the fine motor functions (see Methods; **Fig. 3e**, top panel, Supplementary **Fig. 9b**). The two populations were statistically different at D79 (**Fig. 3e**, top and bottom panels). In adult stroke, the lesion volume is not correlated with the degree of motor impairment (39,40). In order to check whether a correlation between the lesion brain size (**Fig. 3f**, left panel) or brain atrophy (**Fig. 3f**, right panel) and the asymmetry index in the gridwalk test exists, we plotted the data for each stroke animal, and we found no significant correlations. In conclusion, dMCAO in mice models perinatal stroke in terms of brain lesion and motor outcomes. In particular, we did not find any impairment in overall motor performance, and we found an impairment in fine motor outcome across development till adulthood, with no correlation with the extent of the lesion.

**Figure 3.**
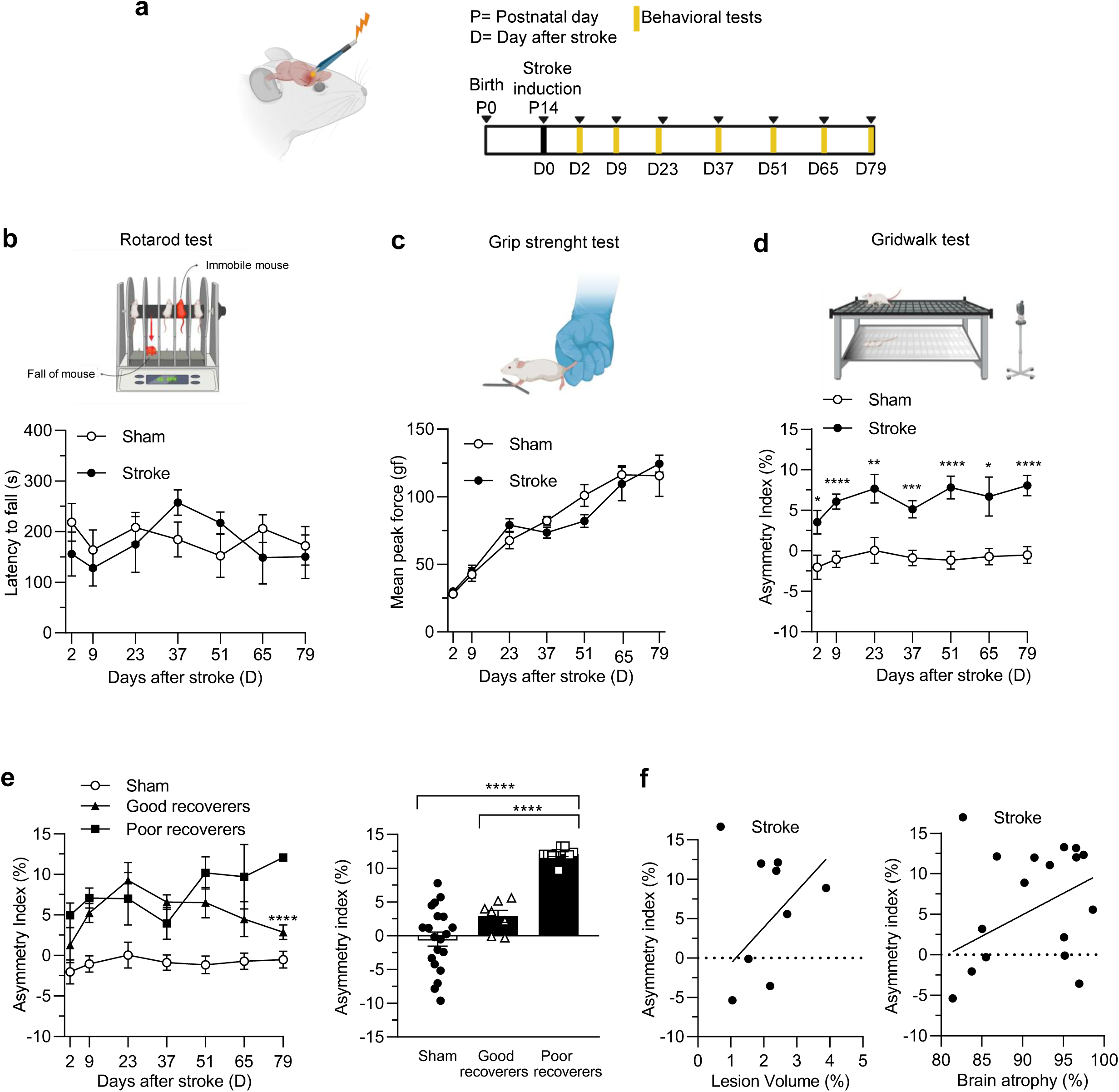
Perinatal stroke impacts fine motor behavior. (**a**) Schematic cartoon of the experimental protocol. (**b**) Top, representative cartoon of the rotarod test. Bottom, quantification of latency to fall in sham (white) and stroke (black) animals showing no significant differences at any point (two-way RM ANOVA; P = 0.9138, 0.9933, 0.9985, 0.5608, 0.8046, 0.9563, 0.9998). (**c**) Representative cartoon of the grip-strength test. Bottom, quantification of peak force in sham (white) and stroke (black) animals showed no significant differences on all the days (two-way RM ANOVA; P > 0.9999, > 0.9999, 0.8821, 0.9698, 0.3266, 0.9957, 0.9603). (**d**) Top, representative cartoon of the grid-walk test. Bottom, quantification of the asymmetry index in sham (white) and stroke animals (black) in which stroke animals showed persistent motor impairments (two-way RM ANOVA, Holm–Sidak; P = 0.0207, P= <0.0001, P= 0.0092, P= 0.0006, P= <0.0001, P=0.0207). (**e**) Top, quantification of the asymmetry index in sham and in “good” and “poor” recoverers across days, and (bottom) specifically at D79 seeing a reduction of the asymmetry index in the good recoverers compared to the poor ones (two-way RM ANOVA, Holm–Sidak; P < 0.0001). (**f**) Correlation between (left) lesion volume and asymmetry index; (P = 0.1653, R²= 0,2937) and (right) brain atrophy with asymmetry index; (P = 0.0856, R²= 0,1963) (All the statistics in the supplementary). (See Methods for details on animal numbers). All histograms show mean ± s.e.m. Statistical significance *P<0.05, **P<0.01, ***P<0.001.

### Perinatal ischemic stroke impacts cognition

Perinatal ischemic stroke in humans is associated with cognitive impairments (41). To assess whether dMCAO in mice results in cognitive deficits, we assessed working memory by performing the spontaneous alternation test in a Y-maze setup (see Methods) (42) based on the high exploration tendency of mice (43). Mice were tested from D2 to D79 (**Fig. 4a**). We found no significant differences between sham and stroke mice in terms of either spontaneous alternation or the number of entries (**Fig. 4b**). Since perinatal stroke is associated with spatial memory deficits (44), as an adult stroke (45), we studied spatial-novelty memory (42) after perinatal-induced dMCAO by performing the forced alternation test (42), at D9, D37, and D79 (**Fig. 4c**). Specifically, we quantified spatial novelty by determining two measures, the percentage of forced alternation, and the percentage of time spent in the novel arm (%). We found that fully-adult (D79) stroke animals spent less time in the novel arm in comparison to sham (**Fig. 3c**, right panel), but no statistical difference in the forced-alternation index (**Fig. 4c**, left panel), suggesting that perinatal ischemic stroke impacts spatial novelty memory.

**Figure 4.**
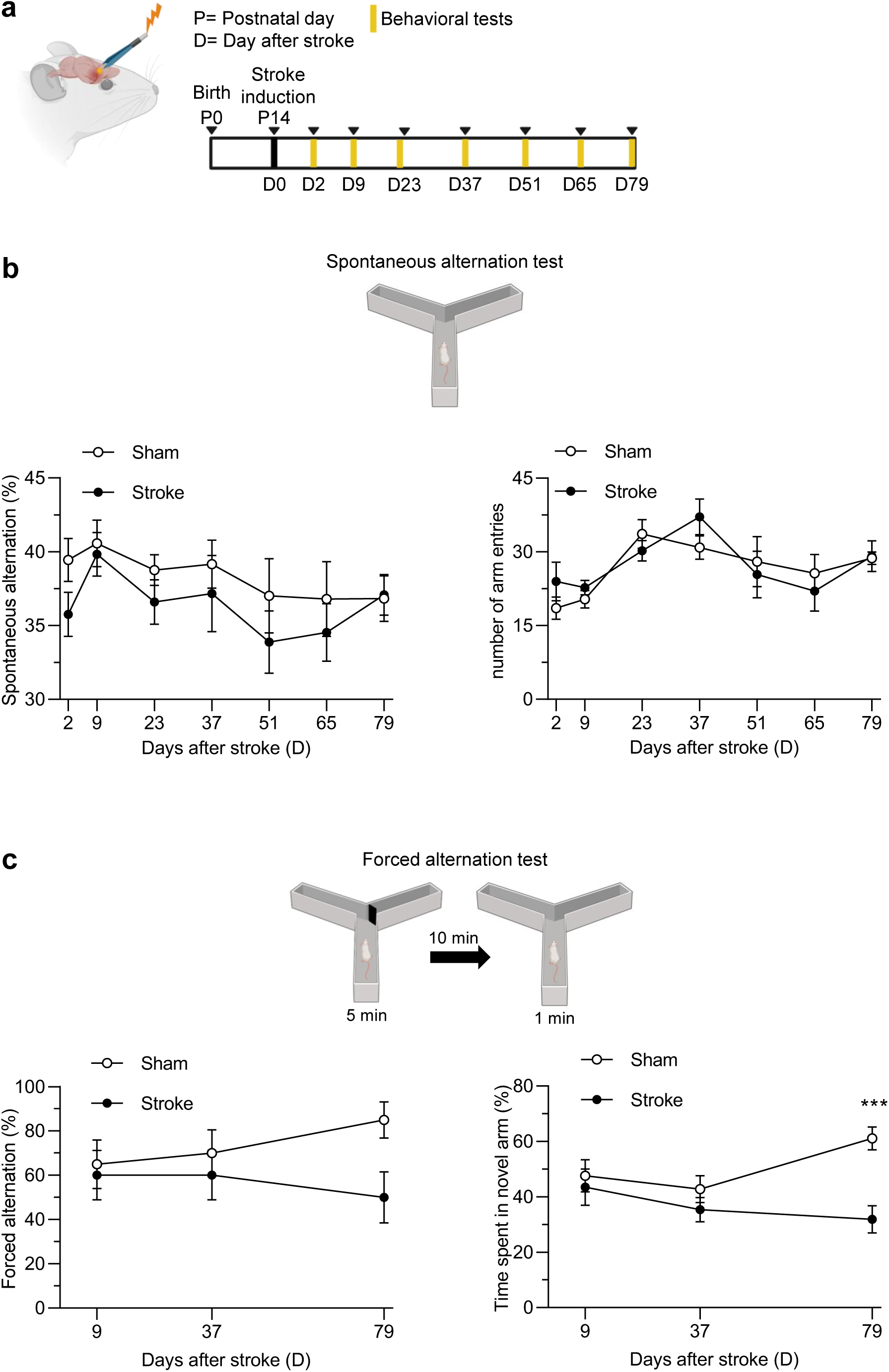
**Perinatal stroke impacts cognitive functions**. (**a**) Schematic cartoon of the experimental protocol. (**b**) Top, representative cartoon of the spontaneous alternation test. Bottom left, quantification of spontaneous alternation across days in sham (white) and stroke (black) animals showing no differences between groups across testing days (two-way RM ANOVA, Holm–Sidak; P = 0.4822, >0.9999, 0.8735, 0.9946, 0.9546, 0.9913, >0.9999), and (bottom right) for the quantification of the number of arm entries during trials (two-way RM ANOVA, Holm–Sidak; P = 0.8791, 0.9344, 0.9603, 0.7389, 0.9998, 0.9949, >0.9999). (**c**) Top, representative cartoon of the forced alternation test. Bottom left: quantification of forced alternation in sham (white) and stroke (black) animals showing no significant differences across days for the percentage of forced alternation (two-way RM ANOVA, post hoc Holm–Sidak, P = 0.9847, 0.8892, 0.0532) while (bottom right) significantly less time in the novel arm at the last testing day was detected in the stroke animals (two-way RM ANOVA, Holm–Sidak; P = 0.955, 0.5973, P = 0.0002). (See Methods for details on animal numbers). All histograms depict mean ± s.e.m. Statistical significance *P<0.05, **P<0.01, ***P<0.001.

### Perinatal ischemic stroke affects the structure of skeletal muscles

In both human and animal studies, adult stroke is correlated with modifications in paretic muscles (46), and with a reduction of motility, motor endurance, and tensile strength (47). However, evidence on the impact of perinatal stroke on skeletal muscle structure is limited (4). To assess whether dMCAO impacted the structure of muscles serving the contralateral upper limb, at the end of the behavioral motor assessment (D79), the contralateral extensor digitorum muscles were isolated, and the sarcomers’ length was measured (**Fig. 5a**). We found that the length of sarcomeres, essential to anchor points for the thin actin filaments (48), was significantly shorter in stroke animals (1.3 ± 0.1 µm), compared to sham (1.5 ± 0.1 µm) (**Fig. 5b**, Supplementary **Fig. 9c**). Furthermore, to assess muscle damage, we analysed hematoxylin/eosin (H/E) stainings (**Fig. 5c**), and found tissue damages in terms of fiber overstaining and fiber vacuolization (**Fig. 5d**) (49). Sarcomere length and damage index did not significantly correlate with fine motor performance (asymmetry index) or brain lesion measures (brain atrophy and lesion volume) in sham and stroke animals (**Fig. 5e**). Taken together, our data show that perinatal ischemic stroke alters the structure of skeletal muscle in the contralateral limb, with no detectable correlation with performance on the gridwalk test (asymmetry index) or with brain lesion volume.

**Figure 5.**
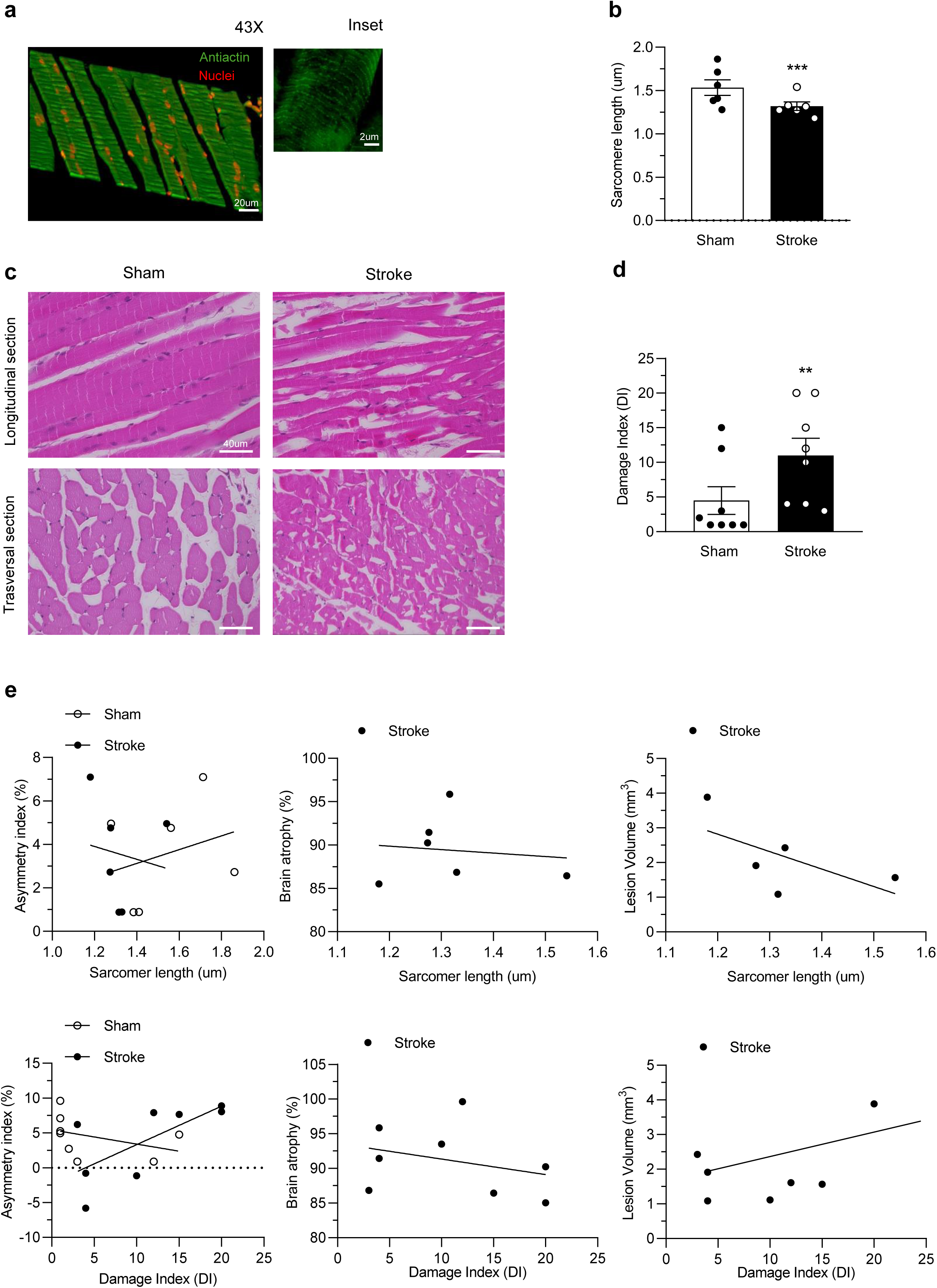
Perinatal stroke affects the structure of skeletal muscles. (**a**) Representative histological section of the extensor digitorum muscle 79 days after perinatal stroke stained with an anti-actin antibody (43× magnification; scale bar, 20 μm). Inset: single sarcomere (scale bar, 2 μm). (**b**) Quantification of sarcomere length in sham (white) and stroke (black) animals showing a reduction in the stroke animals (Mann-Whitney test; P=0.0006). (**c**) Representative hematoxylin/eosin-stained sections of skeletal muscle from sham and stroke mice 79 days after perinatal stroke. Scale bars of 40 μm are labeled only in the first image, while the following images display unlabeled scale bars. (**d**) Quantification of the damage index (combined fiber overstaining and vacuolization score) in sham (white) and stroke (black) animals, showing an increase of this damage index in the stroke animals (Mann-Whitney test; P =0.0208). (**e**) No correlations were found between muscle modification and stroke impairments. Top left, correlation between sarcomere length and gridwalk asymmetry index in sham (white) and stroke (black) animals (R² = 0.07895, P = 0.5896 for sham; R² = 0.02058, P = 0.7863 for stroke). Top center, correlation between sarcomere length and brain atrophy in stroke animals (R² = 0.01501, P = 0.8171). Top right, correlation between damage index and lesion volume in stroke animals (R² = 0.387, P = 0.2625). Bottom left, correlation between damage index and gridwalk asymmetry index in sham (white) and stroke (black) animals (R² = 0.1488, P = 0.3453 for sham; R² = 0.4744, P = 0.0589 for stroke). Bottom center, correlation between damage index and brain atrophy in stroke animals (R² = 0.09621, P = 0.4546). Bottom right, correlation between damage index and lesion volume in stroke animals (R² = 0.03294, P = 0.5933). (See Methods for details on animal numbers). All histograms depict mean ± s.e.m., and circles represent individual animals. Statistical significance *P<0.05, **P<0.01, ***P<0.001.

### Perinatal ischemic stroke affects the gut microbiota bacteria population

To study the gut microbiota composition at the species level (50), we collected feces samples from stroke or sham mice at D23 after surgery and analysed the 16S Ribosomal RNA gene. Our analysis revealed no significant differences between stroke and sham animals for the relative abundance at the Phylum level (**Fig. 6a**). Both stroke and sham microbiota profiles were dominated by Firmicutes (∼88%) followed by Bacteroidota (10%). Stroke samples harbored slightly higher abundances of Deferribacterota (1.15% and 0.07%), Proteobacteria (0.39% and 0.18%), while higher abundances of Verrucomicrobiota were detected in fecal samples derived from sham mice (0.91% and 0.03%, respectively) (**Fig. 6a**). At the species level, we quantified the alpha and beta diversity, indicating how bacterial populations vary within or between samples (51), respectively. We did not observe a statistical difference in the alpha diversity, in terms of Shannon and Simpson indexes (**Fig. 6b**) (52), nor in the beta diversity (**Fig. 6c**). However, the differential abundant analysis (DAA) identified a total of 41 bacterial species with significantly different distribution between stroke and sham mice. Of these, 28 species showed non-zero log-fold changes (LFC), of which 24 were enriched in stroke mice (positive LFC values; **Fig. 6d**, in red) while 4 species were depleted (negative LFC; **Fig. 6d**, in blue), revealing an increase in Gram-positive bacteria population (most of them belonging to the Lachnospiraceae and Oscillospiraceae families). Our data show that perinatal ischemic stroke alters the bacterial populations within the gut microbiota.

**Figure 6.**
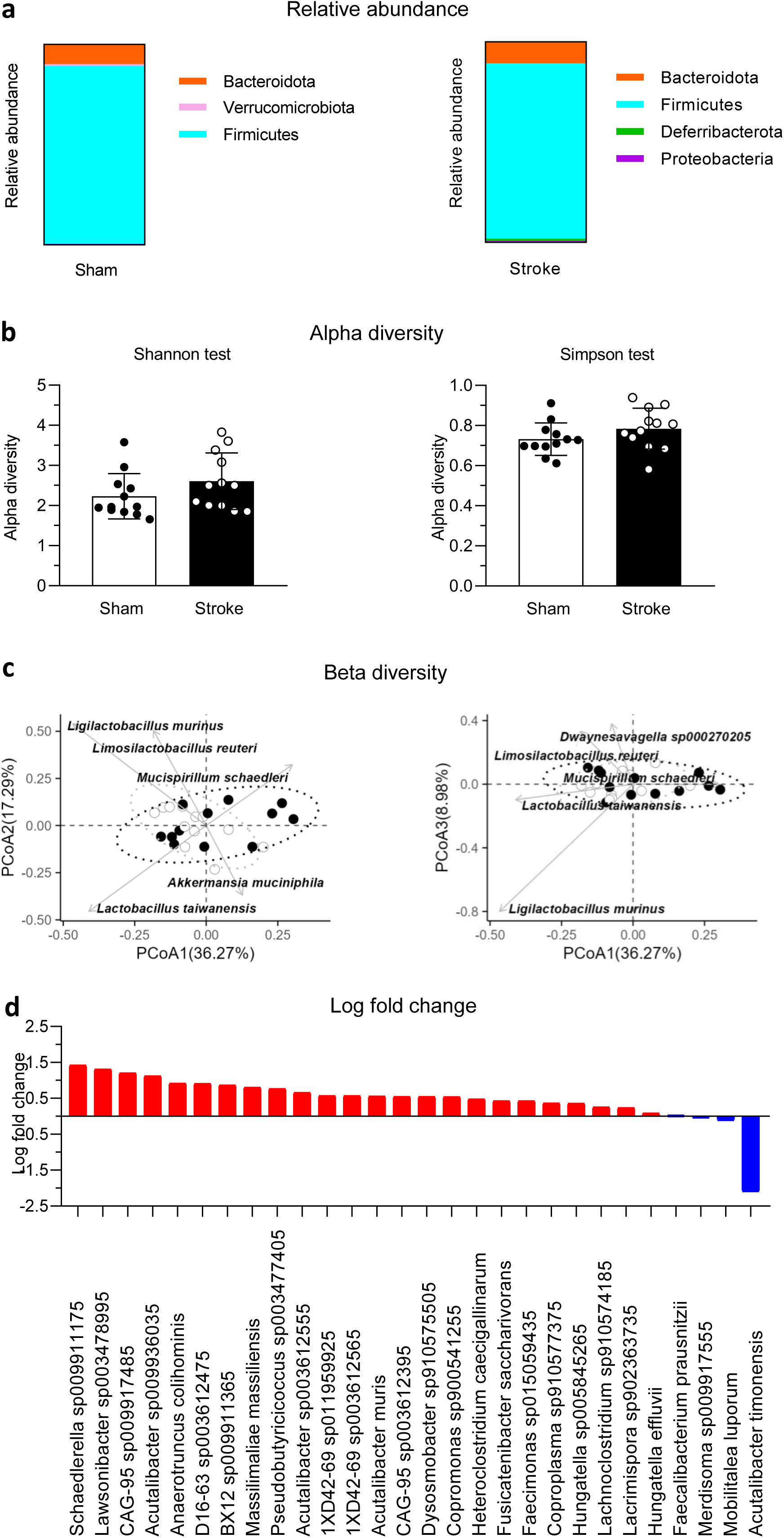
Perinatal stroke affects the gut microbiota bacteria population. (**a**) Relative abundance of bacterial phyla in sham and stroke animals assessed by 16S rRNA V1–V4 sequencing revealed no significant differences. (**b**) Quantification of α-diversity in sham (white) and stroke (black) animals. Shannon index (Student’s t-test; P = 0.4671) and Simpson index (Student’s t-test; P = 0.1794) showing no significant differences. (**c**) Quantification of β-diversity in sham (white) and stroke (black) animals, showing no differences between the two groups (PERMANOVA P = 0.385; R^2^ = 0.04459). (**d**) Log fold change (LFC) analysis showing bacterial taxa increased (red) or decreased (blue) in stroke compared with sham animals. All histograms depict mean ± s.e.m., and circles represent individual animals. (See Methods for details on animal numbers). Statistical significance *P<0.05, **P<0.01, ***P<0.001.

### Fecal microbiota transplantation reduces lesion volume after perinatal stroke

To determine whether FMT exerted similar effects as shown against pathological hallmarks in several pathologies, both in humans and animal models (22,53); in our perinatal stroke model, we performed FMT in stroke mice a few days after stroke induction. Fresh fecal samples from naïve mice were used for transplantation. After MCAO, all mice received oral streptomycin for two days, followed by fecal supernatant gavage in the FMT groups on days 3 and 4 as previously described (Spychala et al. 2018) (**Fig. 7a**). We quantified lesion volumes at D79 in vehicle-treated and FMT-treated mice. Vehicle-treated animals exhibited a *core* hemispheric lesion of 6.565 ± 0.001 mm³, corresponding to 5.68% of the ipsilateral hemisphere. In contrast, FMT-treated mice showed a reduced lesion volume of 3.544 ± 0.001 mm³ (2.82% of the ipsilateral hemisphere), representing an absolute reduction of 3.020 mm³ (a 2.85% reduction relative to the hemisphere) (**Fig. 7b**). These findings align with previously published data demonstrating the neuroprotective effects of FMT (54). We next evaluated whether FMT could also mitigate brain atrophy. Analysis of both whole-brain and hippocampal atrophy revealed no significant differences between vehicle- and FMT-treated groups (**Fig. 7c,d**). This result is consistent with earlier work reporting no detectable change in brain atrophy following FMT (22).

**Figure 7.**
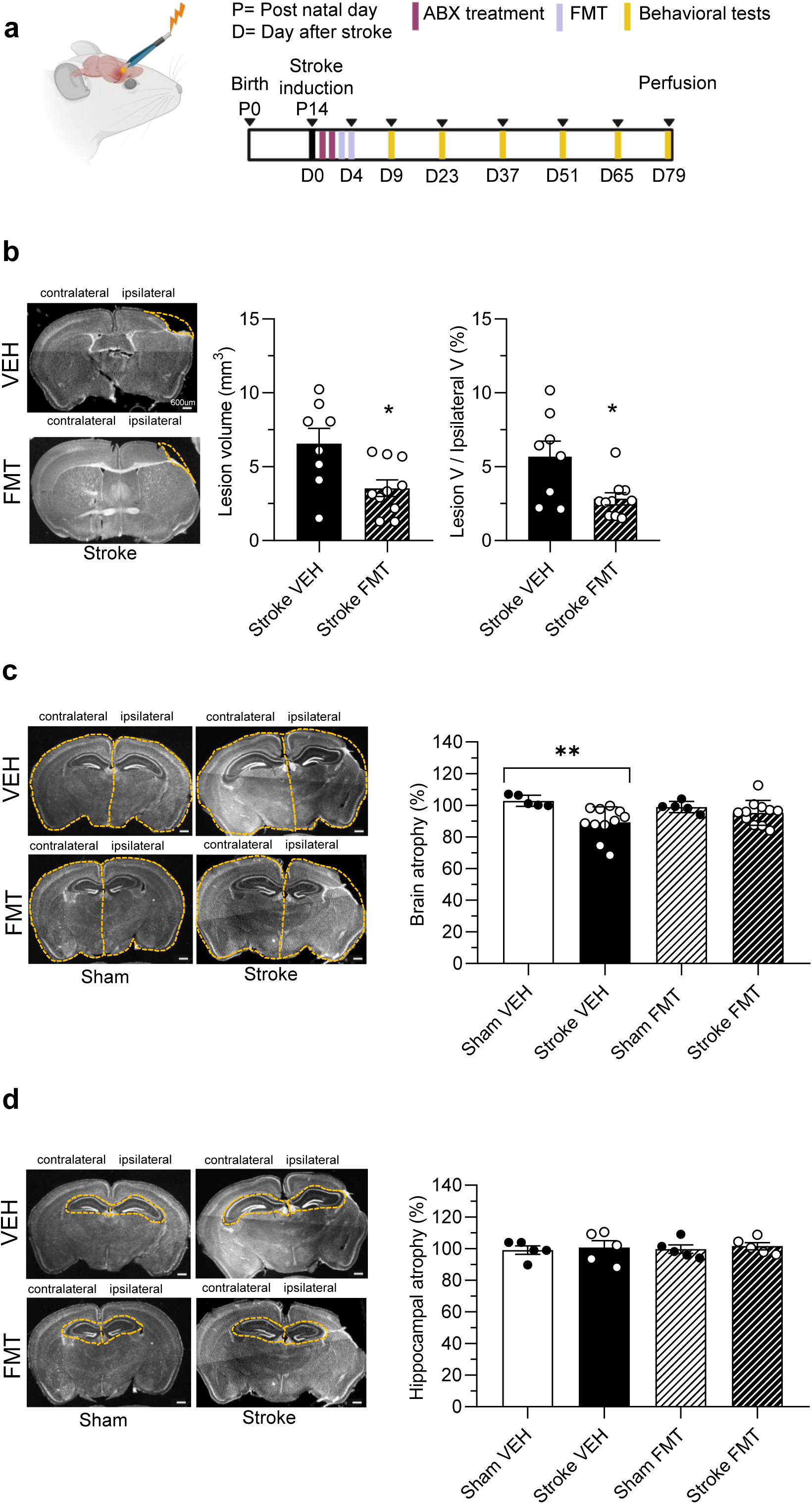
Fecal microbiota transplantation reduces lesion volume after perinatal stroke. (**a**) Schematic cartoon of the experimental protocol. (**b**) Left, representative coronal section of a vehicle mouse stroke brain (top) and FMT stroke mouse brain (bottom) 79 days after perinatal stroke. Right, the lesion volume quantification in stroke animals with the percentage of the lesion compared to the ipsilateral hemisphere in stroke VEH, and stroke FMT; FMT significantly reduced lesion volume compared to vehicle (Student’s t-test; P = 0.0152) and the related percentage of lesion to the ipsilateral hemisphere (Student’s t-test; P = 0.014). (**c**) Left, representative coronal sections of the ipsilateral and contralateral hemispheres of a sham vehicle and FMT mouse brain (left) and stroke vehicles and FMT mouse brain (right) 79 days after perinatal stroke. Right, brain atrophy quantification. A significant difference was observed between sham VEH and stroke VEH groups (two-way RM ANOVA, Holm–Sidak *P < 0.05). No significant differences were observed among the other groups (all statistics in the supplementary). (**d**) Left, representative coronal sections of the ipsilateral and contralateral hippocampus of a sham vehicle and FMT mouse brain (left) and stroke vehicles and FMT mouse brain (right) 79 days after perinatal stroke. Right, quantification of hippocampal atrophy showing no significant differences between groups (two-way RM ANOVA, Holm–Sidak P = 0.342). (all statistics in the supplementary). (See Methods for details on animal numbers). All the D79 images have a 600 μm scale bar. The scale bar is labeled only in the first image, while the following images display unlabeled scale bars. Data shown as mean ± SEM. Statistical significance *P<0.05, **P<0.01, ***P<0.001.

### Fecal microbiota transplantation improves motor behavior after perinatal stroke

Few studies showed a recovery of functions following FMT treatment to a certain extent after a stroke (55). To assess whether restoring gut microbiota balance could improve also motor deficits, we performed the gridwalk test and evaluated the asymmetry index across all the developmental ages tested (D2-D79), (**Fig. 8a**). Interestingly, stroke mice treated with FMT showed an improvement at D79 in the motor performance, shown as a significant reduction in asymmetry similar to sham mice, in comparison to stroke mice treated with vehicle (**Fig. 8b**, left panel). To assess whether FMT treatment led to overall improvement during development, we analyzed the area under the curve (AUC) of the gridwalk test across seven time points. We did not find a significant effect of FMT treatment (**Fig. 8b**, right panel), suggesting that improvements in motor performance appear only later in adulthood. In order to assess whether FMT recover also cognitive function in the spatial novelty, we performed in the same groups of mice the forced alternation test at D9, D37, and D79, and we analyzed the percentage of alternation (**Fig. 8c**, top panels) and the time spent in the novel arm (**Fig. 8c**, bottom panels), including the AUC. No significant differences were observed between groups, indicating that the FMT does not ameliorate the deficit in spatial novelty in stroke mice. Taken together, our data show that FMT in stroke mice specifically recovers fine motor deficits.

**Figure 8.**
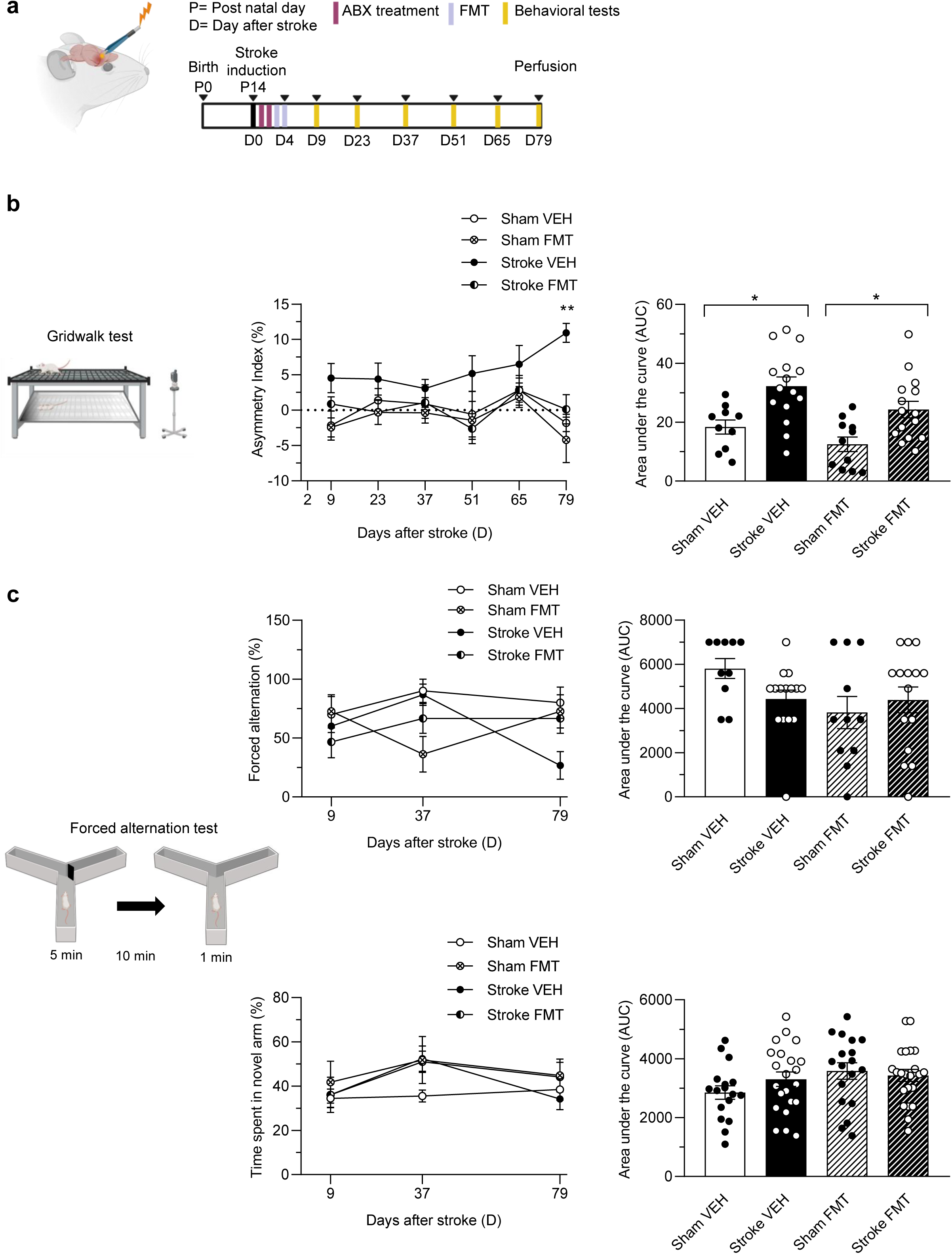
Fecal microbiota transplantation improves motor behavior after perinatal stroke. (**a**) Schematic cartoon of the experimental protocol. (**b**) Left, representative illustration of the gridwalk test. Center, quantification of the asymmetry index in Sham Vehicle (white), Stroke Vehicle (black), Sham FMT (white with “X”), and Stroke FMT (half-black/half-white) groups across days. Significant differences were detected at D79, where stroke FMT animals performed better than stroke vehicles (P = 0.0016), and stroke vehicles differed significantly from both Sham FMT (P = 0.0237) and Sham vehicles (P < 0.0001) (Two-way RM ANOVA, Holm–Sidak). Right, area under the curve (AUC) for gridwalk asymmetry for Sham VEH, Stroke VEH, Sham FMT, and Stroke FMT; Stroke VEH animals showed higher impairment than Sham VEH (P = 0.0093), while Stroke vehicles differed significantly from Sham FMT (P < 0.0001) and Sham FMT differed from Stroke FMT (P = 0.0278) (Two-way RM ANOVA; Tukey). (**c**) Left, representative illustration of the forced alternation test. Top center, quantification of forced alternation performance in Sham VEH, Stroke VEH, Sham FMT, and Stroke FMT across days. A significant group difference emerged at D79, where stroke vehicles performed worse than sham VEH (P = 0.0415) (Two-way RM ANOVA; Holm–Sidak). Top right, AUC quantification of forced alternation performance for sham VEH, stroke VEH, sham FMT, and stroke FMT. No significant differences were detected (Tukey, all P > 0.1). Bottom center: quantification of time spent in the novel arm for the four treatment groups across days. No comparisons reached significance across any of the timepoints (Two-way RM ANOVA; Holm–Sidak. Bottom right, AUC quantification of time spent in the novel arm. No significant differences were detected across groups (Holm–Sidak, all P > 0.24). (See Methods for details on animal numbers). All data are presented as mean ± SEM, with points representing individual animals. Statistical significance *P<0.05, **P<0.01, ***P<0.001.

## DISCUSSION

Perinatal ischemic stroke is a brain injury occurring early in development and is the main cause of cerebral palsy in survivors (1,2). The impact of perinatal stroke, both within the brain and in peripheral systems, remains poorly understood compared with adult stroke, likely due to the limited number of available preclinical models (24). A deeper understanding of perinatal ischemic stroke is important to open up new potential therapeutic approaches (56). In this study, we adapted the dMCAO model, widely used in adult stroke research (26), to the perinatal age on the CD1 mouse strain. Previous work validated this approach at P12 in SCID CB-17 mice (57). However, performing dMCAO at P14 increased post-operative survival and enabled longitudinal behavioral assessment, while remaining within a developmentally relevant window corresponding to human perinatal period (24). Improved survival compared with earlier induction ages is consistent with prior reports (58,59), allowing for subsequent behavioral testing. The lesion pattern induced by dMCAO at P14 was localized primarily to the somatosensory cortex, consistent with adult dMCAO studies (60,61) and with clinical imaging data in perinatal stroke, where infarcts typically involve 5-15% of the cerebral hemisphere (62,63). Progressive lesion expansion between acute and chronic phase, and brain atrophy observed at late time point are in line with evidence that secondary injury mechanisms, including inflammation and edema, contribute to delayed tissue loss following stroke (64,65). While variability in lesion volume across perinatal stroke models has been widely reported (23,66), our study is, to our knowledge, the first to longitudinally characterize both global brain and hippocampal atrophy following perinatal dMCAO. At the cellular level, we observed a progressive reduction in neuronal density from the acute to chronic phase, which is consistent with the absence of detectable brain atrophy in the acute phase and its presence later on. This likely reflects delayed neuronal loss in the penumbral region, which may not yet be as evident during the early post-injury period compared to the chronic period. These findings further support the contribution of secondary injury processes to progressive tissue degeneration after stroke. We observed increased CXCR4 expression during the acute phase, followed by a reduction in the perilesional cortex at adulthood. This temporal pattern is consistent with transient CXCR4 upregulation shortly after adult ischemic stroke (67). Evidence that the CXCR4-CXCL12 signaling axis promotes recovery after adult ischemic stroke by enhancing neurogenesis, angiogenesis, and progenitor cell recruitment to injured brain regions (8,67–69), and suggests that CXCR4 may play a role in functional recovery also after a perinatal injury, a topic that we plan to address in the future. Functionally, our data indicate persistent fine motor impairment following perinatal stroke, with relatively preserved gross motor function. This dissociation closely mirrors clinical observations in children with unilateral cerebral palsy (30) (70). The absence of deficits in grip strength and rotarod performance is consistent with literature reporting limited sensitivity of these tests beyond acute and subacute phases, particularly in developing or chronic stroke models (71–73). In contrast, the gridwalk test reliably detected fine motor deficits from the early post-stroke period through adulthood, consistent with its established sensitivity in adult stroke models (74,75). Our longitudinal design enabled assessment of fine motor outcomes from the early post-injury period through adulthood, extending previous perinatal stroke studies that focused on isolated developmental stages. The dissociation between persistent fine motor deficits and preserved gross motor function suggests selective vulnerability of corticospinal and sensorimotor integration circuits. Fine motor tasks rely on precise cortical control, interhemispheric coordination, and intact somatosensory feedback, functions that are disproportionately affected by focal cortical injury and disrupted network connectivity (76). In contrast, gross motor behaviors are supported by more distributed and redundant motor pathways, including subcortical and spinal circuits, which may compensate more effectively during development (77). This pattern closely parallels clinical observations in unilateral cerebral palsy and supports the idea that perinatal stroke preferentially disrupts high-resolution motor control rather than global motor capacity. Cognitive assessment revealed selective impairment in spatial novelty, with preserved working memory. This pattern aligns with adult stroke studies using the Y-maze that report intact working memory in the absence of hippocampal damage (78,79), while impairments in spatial memory have been documented using forced alternation paradigms (42),(80). The absence of working memory deficits despite impaired spatial novelty processing points to differential susceptibility of cognitive networks. Working memory relies on distributed fronto-hippocampal and prefrontal circuits that may remain functionally intact in the absence of direct hippocampal damage(81). In contrast, spatial novelty detection and exploratory behavior are more sensitive to subtle alterations in cortical-hippocampal communication and network integration (82). Furthermore, the lack of correlation between behavioral outcomes and lesion volume or atrophy is consistent with reports suggesting that post-stroke deficits depend more on network disruption and altered connectivity than on lesion size alone (57,83). However, other studies have reported no differences (84,85). Moreover, the stratification of stroke animals into good and poor recoverers is consistent with findings from both experimental and clinical stroke studies, further supporting the translational relevance of this perinatal model, as most recovery studies have focused on adult stroke (86,87). Beyond central outcomes, our study provides novel evidence that perinatal stroke induces long-term peripheral alterations in skeletal muscle. Structural changes observed in the extensor digitorum muscle, including reduced sarcomere length, increased vacuolization, and fiber damage, are consistent with muscle pathology described following adult stroke (88,89), and other neuromuscular conditions (90). The use of a quantitative damage index enabled objective assessment of muscle degeneration and revealed strong concordance with immunohistochemical markers of post-stroke muscle pathology (88,89). These findings extend existing knowledge by demonstrating that early-life brain injury has lasting downstream effects on muscle structure, potentially contributing to persistent motor deficits. Several, not mutually exclusive, mechanisms may underlie this phenomenon. First, reduced neural drive to affected muscles may lead to activity-dependent disuse atrophy, impairing sarcomere addition during postnatal growth. Second, perinatal stroke may create a developmental mismatch between muscle growth and motor neuron maturation, resulting in aberrant muscle architecture that persists into adulthood. Third, stroke-induced neuroinflammation and systemic inflammatory signaling may directly disrupt myogenesis and sarcomere organization during critical developmental windows. These mechanisms are particularly relevant in early life, when muscle structure is highly plastic and dependent on coordinated neural and metabolic cues (91). We further explored gut microbiota alterations following perinatal stroke, an area that remains poorly characterized (92). In contrast to adult stroke models reporting reduced diversity and broad compositional shifts (93,94) we did not observe changes in α- or β-diversity or phylum-level abundance. Instead, we detected selective increases in gram-positive taxa, particularly within the Lachnospiraceae and Oscillospiraceae families, which are associated with anti-inflammatory functions and metabolic and neurological health (95,96). These findings suggest a compensatory microbiota response to post-stroke inflammation rather than overt dysbiosis, potentially reflecting developmental differences in gut–brain interactions. The concurrent presence of muscle pathology and gut microbiota alterations raises the possibility of a bidirectional gut–muscle axis contributing to post-stroke outcomes. A speculative but compelling model is that perinatal stroke leads to reduced mobility and altered feeding or autonomic regulation, which in turn modifies gut microbial composition. Even subtle shifts in specific taxa may reduce short-chain fatty acid availability or promote low-grade systemic inflammation, thereby impairing muscle repair, regeneration, and metabolic efficiency. Impaired muscle health could then exacerbate motor deficits, further reinforcing changes in gut microbiota through reduced physical activity, forming a self-sustaining pathological loop linking brain injury, gut dysregulation, and muscle dysfunction. Clinical studies have linked gut microbial diversity to skeletal muscle mass, strength, and quality after stroke (97), supporting this interpretation. Finally, we investigated FMT as a therapeutic strategy. FMT reduced infarct size and improved fine motor performance, consistent with reports in adult stroke and other neurological disorders (22,55,98), while cognitive outcomes remained unchanged, as previously observed (99). This selective motor improvement may reflect the involvement of the motor cortex *penumbra* and its capacity for plasticity. The selective efficacy of FMT on motor, but not cognitive, outcomes further supports the existence of differential therapeutic windows and circuit-specific responsiveness. One possibility is that the motor cortex penumbra retains greater plasticity and remains more responsive to anti-inflammatory and metabolic modulation during early development. In contrast, cognitive networks may require longer treatment duration or additional circuit-specific signals to achieve functional recovery. Restoring microbial balance early after stroke may therefore mitigate neuroinflammation and support motor recovery, particularly during development (100–102). However, moving toward clinical trials in neonates will require careful, stepwise validation. Key preclinical steps include defining the optimal therapeutic window, donor microbiota composition, dosing strategies, and durability of effects, as well as establishing causal links between specific microbial taxa, immune modulation, and neuroplasticity. Collectively, this work substantially reshapes our understanding of perinatal ischemic stroke, shifting the conceptual framework from a disorder confined to the central nervous system toward a systemic condition involving coordinated alterations across the brain-gut-muscle axis. Rather than viewing motor and cognitive impairments solely as direct consequences of focal brain injury, our findings indicate that early-life stroke initiates long-lasting peripheral adaptations that may actively contribute to functional outcomes. This paradigm shift expands the scope of perinatal stroke research and identifies new biological targets beyond the injured brain itself. Several limitations should be considered. Behavioral analyses were affected by variable sample sizes, as not all animals reliably engaged in each task, reducing statistical power and potentially introducing selection bias. Behavioral performance is also influenced by non-neurological factors such as motivation, stress reactivity, and handling, which may confound the interpretation of stroke-related deficits. In addition, exposure to isoflurane anesthesia during early postnatal development may itself affect neurodevelopment and long-term behavior, potentially interacting with ischemic injury. Perinatal stroke models further exhibit inherent variability in lesion size and location, contributing to heterogeneity in outcomes. Finally, differences in brain development and environmental complexity between rodents and humans limit direct translation of these findings to clinical neonatal stroke. In summary, this study establishes the P14 dMCAO model as a robust and translationally relevant platform for investigating perinatal ischemic stroke while fundamentally expanding the conceptual boundaries of the field. By integrating longitudinal behavioral outcomes with analyses of brain structure, skeletal muscle pathology, gut microbiota composition, and microbiome-based intervention, our work underscores the importance of systemic interactions in shaping long-term recovery after early developmental brain injury. This integrative framework opens new avenues for therapeutic development and suggests that targeting peripheral systems may be essential for improving outcomes in perinatal stroke and potentially other pediatric neurological disorders.

## METHODS

### Sex as a biological variable

Our study included both male and female subjects. No sex-dependent differences were observed, and the findings were comparable between sexes.

### Animals

CD1 mice were housed in the Animal Facility at IRP under controlled conditions (22°C, 12 h light/dark cycle) and had ad libitum access to standard pellet chow and reverse osmosis water. All surgical procedures were conducted under deep anesthesia, and all the animal manipulations were made keeping the maximum attention to minimize the animal suffering. In accordance with study design considerations for sex as a biological variable, both male and female mice were included in all experiments (all the animals described in **Table 1**). Animals were distributed across experimental groups without sex-based exclusion. Data were analyzed to assess potential sex-dependent effects, and no significant differences were observed between male and female mice for the measured outcomes. These findings indicate that the reported results are consistent across both sexes within the conditions tested.

**Table 1.** Animals Included in Experimental Procedures. This table summarizes the number of animals used in the experimental procedures.

| **Procedures** | **Day** | **Sham** | **Stroke** | **Sham**<br>**vehicles** | **Sham**<br>**FMT** | **Stroke**<br>**vehicles** | **Stroke**<br>**FMT** |
| --- | --- | --- | --- | --- | --- | --- | --- |
| Lesion volume | D2 |  | 3 |  |  |  |  |
|  | D79 |  | 8 |  |  | 8 | 10 |
| Brain atrophy | D2 | 4 | 3 |  |  |  |  |
|  | D79 | 16 | 15 | 5 | 5 | 11 | 11 |
| Hippocampal<br>atrophy | D2 | 3 | 3 |  |  |  |  |
|  | D79 | 16 | 15 | 5 | 5 | 5 | 5 |
| Rotarod test | D2 – D79 | 8 | 8 |  |  |  |  |
| Grip strength test | D2 – D79 | 8 | 8 |  |  |  |  |
| Gridwalk test | D2 | 20 | 23 |  |  |  |  |
|  | D9 | 23 | 24 | 14 | 14 | 14 | 15 |
|  | D23 | 19 | 17 | 15 | 12 | 15 | 15 |
|  | D37 | 24 | 21 | 13 | 13 | 11 | 15 |
|  | D51 | 23 | 22 | 13 | 13 | 12 | 13 |
|  | D65 | 19 | 15 | 13 | 12 | 11 | 11 |
|  | D79 | 20 | 16 | 12 | 9 | 11 | 13 |
| Spontaneous<br>alteration test | D2 – D79 | 12 | 12 |  |  |  |  |
| Forced alternation<br>test | D2 | 16 | 16 |  |  |  |  |
|  | D37 | 16 | 16 | 10 | 11 | 15 | 15 |
|  | D79 | 16 | 16 | 10 | 11 | 15 | 15 |
| Sarcomere analysis | D79 | 6 | 6 |  |  |  |  |
| DI analysis | D79 | 8 | 8 |  |  |  |  |
| CXCR4 analysis | D2 | 3 | 3 |  |  |  |  |
|  | D79 | 7 | 7 |  |  |  |  |
| Microbiota analysis | D23 | 12 | 12 |  |  |  |  |

### Distal middle cerebral artery occlusion (dMCAO)

dMCAO was performed as previously described (26) and adapted at P14. Mice were randomly divided into two different groups: the control (sham) group and the stroke group. Mice were anesthetized with isoflurane (4% for induction, 1.5–2.5% for maintenance) at a flow rate of 1 L/min. Prior to surgery, a combination of analgesics, dexamethasone (2 mg/kg, Dexadreson) and carprofen (5 mg/kg, Rymadil) was administered via intramuscular and intraperitoneal injection, respectively, and the body temperature was maintained at ∼36°C using a heating pad (KF Technology, Italy) throughout the procedure. Once the mice were unresponsive to tail and paw reflexes, they were positioned in an apparatus for dMCAO surgery. The area between the ear and eye was shaved, disinfected, and immobilized using medical adhesive tape in preparation for incisions. A horizontal incision was made between the eye and ear canal, followed by a vertical incision from the ear canal, creating an “L”-shaped skin flap. The same incisions were made on the right temporalis muscle to expose the skull. A craniotomy was then performed below the MCA decussation, creating a rectangular opening without dura, allowing MCA occlusion using 0.25 mm bipolar forceps (GIMA, Italy) connected to an electrosurgical unit (Diatermo 122, GIMA, Italy). If spontaneous recanalization occurred, electrocoagulation was repeated. Finally, the temporalis muscle and skin were repositioned and secured using veterinary glue (Vetbond, 3M, USA). The wound was sealed by gluing the skin flaps together. Sham animals underwent the same procedure but without craniotomy or MCA electrocoagulation (23).

### Fecal microbiota transplantation

Fresh fecal samples were collected from naïve mice and homogenized in NaCl (0.9%) (120 mg feces/1 mL buffer) and then homogenized for 5 minutes until a paste-like consistency was achieved, as described by (22,103). The supernatant was transferred into new tubes and used for transplantation. Sham or stroke mice received oral gavage administration of antibiotic streptomycin sulfate salt (500 mg/Kg, Sigma-Aldrich) in 50 μL sterile (ABX) for the first two days after the MCAO. On days 3 and 4 after stroke, 40 μL of the fecal supernatant was administered by oral gavage to the FMT group, while the vehicle group received only NaCl (0.9%).

### Behavioral tests

After the dMCAO surgery, the sham and stroke animals and the sham vehicle and FMT and stroke vehicle and FMT underwent behavioral testing from the D2, D9, and every two weeks until the D79. On the day of behavioral testing, mice were placed in the experimental room 30 minutes prior to the start of the tests for acclimatization. All equipment was cleaned with 70% ethanol after each trial. Behavioral tests were recorded and analyzed in blind using customized MATLAB scripts (MathWorks, Massachusetts, USA).

#### Rotarod test

To evaluate general motor coordination and balance, the rotarod test was performed using the rotarod apparatus (Mouse Rota-Rod, Ugo Basile Biological Research Apparatus, Comerio, Italy) as described by (13). The apparatus featured a rotating rod (3 cm in diameter) divided into separate lanes, allowing multiple animals to be tested simultaneously. The rotation speed increased progressively until the mouse fell. Each animal underwent three sessions per day (spaced 30 minutes apart), with the rotation starting at 4 RPM (rotation per minute) and gradually accelerating to a maximum of 40 RPM over 300 seconds. The time to fall (in seconds) was recorded, and the mean latency across the three trials was analyzed for different days of the experimental protocol using a custom MATLAB script. The test was conducted from D2 to D79 after stroke induction.

#### Grip strength test

Forelimb grip strength and overall muscle power were assessed using the Grip Strength Meter (Mouse Grip Strength Meter, Ugo Basile Biological Research Apparatus, Comerio, Italy) (34). The apparatus consisted of a metal bar equipped with a force sensor. Mice were held by the tail and allowed to grasp the metal bar with their forelimbs before being gently pulled backward with consistent force until they released their grip. The force sensor recorded the peak force exerted. Each animal completed three consecutive trials with a 1-minute inter-trial interval. The mean peak force across the three trials was analyzed on different experimental days using a custom MATLAB script. This test was conducted from D2 to D79 after stroke induction.

#### Gridwalk test

Fine motor impairments were assessed using the sensory-motor gridwalk test (13). Mice were allowed to walk freely on an elevated grid (32 × 20 cm) with uniform 11 × 11 mm spaces between the bars, while a camera positioned in front of the apparatus recorded their movement. Each mouse was placed at the center of the grid and allowed to freely walk for 5 minutes. The asymmetry index was calculated manually as the difference between the percentage of left and right foot faults, based on a total of 150 to 200 steps. This analysis was conducted using a custom-designed Graphical User Interface in MATLAB. The test was performed from D2 to D79 following stroke induction. To differentiate between good and poor recovery outcomes, the asymmetry performance of the sham at D79 was used as a reference for healthy motor function. A threshold was defined as two standard deviations from the mean of this control group. This criterion was then applied to the stroke at D79: stroke mice whose motor scores fell within this threshold were classified as having a good recovery outcome; in contrast, those with scores exceeding the threshold were considered poor recoverers, reflecting a level of impairment beyond the natural variability observed in healthy controls and indicating a less favorable post-stroke outcome.

#### Y-maze

To assess cognitive impairments, the Y-maze apparatus (Mouse Y-maze Stoelting) was used (104). The apparatus consisted of three equal arms with a measure of 35cm (arm length) x 5cm (lane width) x 10cm (arm height), with a final wall with a height of 15.5 cm. Two distinct tests were performed: the spontaneous alternation test, used to assess the working memory, in which each mouse underwent a single 10-minute trial, starting at the end of a designated arm, facing the wall and away from the center. The mouse was allowed to explore all three arms freely. An arm entry was recorded when all four paws entered an arm (104,105). The spontaneous alternation percentage was calculated as the number of consecutive entries into three different arms (ABC), divided by the total possible alternations (total arm entries minus two) (42). To prevent odor cues, the maze was cleaned with 70% ethanol after each trial. This test was conducted from D2 to D79 after stroke induction. The second test was the forced alternation test, used to assess the special novelty deficits (42). The test consisted of a 5-minute sample trial (T1),10 minute interval, followed by a 5-minute retrieval trial (T2). In particular, in T1 the mouse was placed in the start arm, facing the wall and away from the center. It was allowed to explore two arms of the Y-maze while access to the third arm was blocked by a sliding door. After 5 minutes, the mouse was returned to its home cage for a 10-minute inter-trial interval. The maze was cleaned with 70% ethanol between T1 and T2 to prevent odor cues. In T2, the block was removed from the previously inaccessible arm, and the mouse was reintroduced into the start arm. It was then allowed to explore all three arms for 2 minutes. Two different parameters were analyzed: first, the time in the novel arm (%), which is defined as the time spent in the novel arm divided by the time spent in all arms during the first minute of the retrieval trial T2. Second, the forced alternation (%), defined as the percent of mice entering the novel arm during T2. The test was performed at D9, D37, and D79 after the stroke induction.

### Tissue preparation and histology

Following completion of all experiments (post D2 and post D79), mice were anesthetized (ketamine 200mg/kg, xylazine 50 mg/kg) and transcardially perfused with 20ml of phosphate-buffered saline (PBS, Sigma-Aldrich, P4417) 1X, followed by 30ml of 4% paraformaldehyde (PFA, in 0.1 M Phosphate Buffer, pH 7.0-7.4) for the brain tissues fixation. Brains were extracted and immersed in 4% PFA at 4°C for post-fixation. The following day, brains were cryopreserved in 30% sucrose solution in 1X PBS before being frozen. Tissue sections (60 µm) were obtained using a vibratome (VT1000S, Leica) and a microtome (Leica SM2010R). For CXCR4 immunostaining, coronal brain slices were quenched for 20 min in 50 mM NH4Cl in PBS, permeabilized, and saturated for 1 h in blocking solution (15% goat serum, 2% BSA, 0.25% gelatin, 0.20% glycine) containing 0.5% Triton X-100. Incubation with primary antibodies was carried out O/N in blocking solution + 0.5% Triton X-100 (anti-Cxcr4 (ab124824, Abcam, Cambridge, UK, 1:200), anti-Neurofilament heavy polypetide (ab4680, Abcam, 1:500), anti-NeuN (MAB377, Sigma-Aldrich, Saint Louis, MO, USA): 1:200, DAPI (D3571, Thermo Fisher Scientific, Waltham, MA, USA): 1:5000). Slices were then washed in PBS and incubated with secondary antibodies diluted in PBS for 2 h. Nuclei were stained with DAPI. For muscle tissue, the extensor digitorum posterior was collected on D79 post-stroke. Samples were fixed by embedding them in OCT, flash-frozen using dry ice, and stored at -80°C. For muscle processing, extensor digitorum muscles were fixed overnight in 4% v/v paraformaldehyde (Sigma, USA) at 4 °C under gentle agitation. Tissues were then dehydrated in an ascending ethanol series (25%, 50%, 70%, 95%, and 100%) before embedding in histology-grade paraffin (Leica biosystems, Germany) using a HistoCore Arcadia H embedding station (Leica biosystems, Germany). Then, embedded samples were sectioned at 5 μm thickness using a fully-motorized rotary microtome Leica RM2265 (Leica biosystems, Germany), collected on superfrost plus adhesion slides (Thermofisher, USA), and stored at room temperature in the dark until histological or immunological staining. Brain lesion volume and brain atrophy were evaluated by staining brain sections (one in every six sections spanning the *core* lesion) with Hoechst 33258 (Thermo Fisher). Sections were then mounted on glass slides with coverslips and Vectashield as mounting medium (Vector Labs). For muscle histochemical and immunohistochemical staining, the sections were deparaffinized in Xylene (Sigma, USA), rehydrated by soaking in a descending ethanol/H_2_O series (100%, 95%, 70%, and 50%), and gently washed under deionized water for 2 minutes. The hematoxylin/eosin (H/E) histological staining was performed following a modified Harris protocol we previously adapted to musculoskeletal tissues (106). Briefly, slides were sequentially stained with H/E, optimizing the contact time with each probe (2 min. and 1 min. for eosin, and hematoxylin, respectively) to obtain balanced visualization and an optimal stain/counterstain ratio. Finally, the sections were washed with running tap water, then mounted with a coverslip and limonene mount medium (Fisher Scientific, United States). For immunostaining, our protocol already optimized for 3D culture, was readapted to be used in skeletal muscle samples (107). Briefly, Rab-Anti-Sarcomeric Alpha Actinin antibody (dilution 1:1000, ab68167, Abcam, USA), and Ms-mMaC myomesin B4 (dilution 1:500 v/v, AB_760349, DSHB, USA) were used as primary unconjugated antibodies. Donkey anti-Rabbit DL488 (dilution 1:50 v/v, Ab96919, Abcam, USA) and Donkey anti-Mouse DL550 (dilution 1:50 v/v, Ab98795, Abcam, USA) were used as secondary conjugated antibodies for a selective visualization. Propidium iodide (dilution 250 µg ml −1, ab14083) was used as nuclear staining. The antigen retrieval was performed using an aqueous citrate buffer at pH 6.0 containing Sodium Citrate dihydrate 0.294% w/v (Sigma, USA), 5 μl Tween 20 (Sigma, USA), and deionized water. Slides were then incubated within a StainTray slide staining system (Merck, Germany) with primary antibodies overnight at 4 °C in the dark. Then, the primary antibody was washed away by rinsing three times in PBS, and samples were incubated with a mixture of secondary antibody and nuclear staining for 30 min at RT in the dark. Finally, slides were mounted with ProLong™ Glass Antifade Mountant (Thermofisher, USA) and stored in the dark at 4 °C.

### Optical and confocal microscopy

All slides were acquired in a random order and in a single session for each litter of animals to minimize errors caused by fluctuation in laser output and degradation of fluorescence. Images for the lesion volume and brain atrophy quantification were acquired using an epifluorescence Leica DM600B Microscope (Leica Microsystems, Germany) at 2.5x magnification. The fluorescence measurements of the Neun and CXCR4 expression were performed using the LSM900 Airyscan 2 confocal microscope (Zeiss, Oberkochen, Germany) equipped with 63x/1.4 Plan-Apochromat oil immersion objective, for the stroke group, 4 images were acquired: 2 from the medium-lateral (ML) and 2 from the peri-lateral (PL) area outside the infarcted zone, one from a superficial and one from a deep section. For the sham group, 2 images were acquired, one from a superficial and one from a deep section. The fluorescence measurements of the muscle tissue were performed using a SP5 confocal microscope (Leica, Germany) equipped with a resonant scanner, using an Objective HC PL APO 40x/1.30 (Leica, Germany) working in oil immersion.

### Lesion volume and brain atrophy quantification

Tissue damage is a critical determinant of disease progression and is commonly assessed through lesion quantification (108). Images were analyzed using Fiji software (109). First, it was manually contoured the ipsilateral hemisphere and infarct area of each session. Lesion volume was calculated as follows:

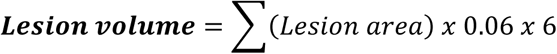

To quantify brain atrophy, the percentage shrinkage of the ipsilateral hemisphere relative to the contralateral hemisphere was calculated. The same brain sections and images used for lesion volume analysis were utilized. Image acquisition was performed using a Leica DM600B Microscope at 2.5× magnification, and measurements were analyzed using Fiji software (109).

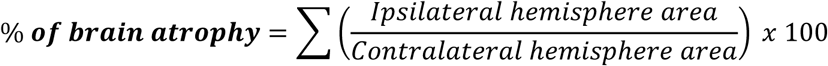

### Damage index for muscle structure

The damage index (DI) was evaluated based on pathological anatomy evidence collected during a deep analysis of H/E stained tissues. Specifically, we evaluated the presence of fiber vacuolization (FV) (in transversal sections) and fiber overstaining (FO) (in longitudinal sections), which are linked to oxidative and ischemic stress, respectively. A score ranging from 0 to 5 was assigned to FV and FO, where 0 = no tissue damage, and 5 = highly damaged tissue. Finally, the DI was calculated according to the following formula: DI = FV x FO. The images for the DI assessment were processed by two independent, skilled scientists following a blind approach, where the tissue condition was unknown during the image analysis.

### Gut microbiota sequencing and analysis

Solid D23 mouse feces were rehydrated in 1 mL of PBS and DTT (0.1% final concentration) and vortexed to ensure homogeneous resuspension. 500µL of rehydrated stools were transferred into a clean tube containing beads and mechanically lysed with a FastPrep-24 5G homogenizer. Total DNA was isolated using the MagMax Microbiome Ultra Nucleic Acid Isolation Kit in combination with KingFisher Flex device (ThermoFisher) following the manufacturer’s instructions. Full-length 16S rRNA sequencing libraries were prepared as described in the publication by Pinzauti et al (110). Sequencing was performed using a GridION Mk1b device (Oxford Nanopore Technologies), enabling real-time, super-accurate base-calling (Quality Threshold Q ≥ 12). Raw fast Q sequencing reads were analyzed using a custom pipeline, inferring taxonomy (110). The resulting taxonomy tables were then imported into RStudio (build v. 524, R v. 4.3.1) for microbial community analysis with the phyloseq tool (v.1.44) (111). Alpha diversity analysis was performed by evaluating the Shannon and Simpson diversity indices. Beta diversity was evaluated by running Principal Coordinate Analysis (PCoA), using Bray-Curtis dissimilarity index and Hellinger standardization method (MicrobiotaProcess v1.14.1) (112).

### Statistical analysis

Statistical analysis was performed with GraphPad Prism 8.0.1. All data were run for a normality test by default in GraphPad Prism 8.0.1. before any statistical comparison tests. Data normally distributed were summarized by the mean ± s.e.m., whereas data non-normally distributed were summarized with a box chart. Pairwise comparisons of quantitative phenotypes between mice of different groups, normally distributed, were assessed by a two-tailed Student’s *t*-test. When more than two groups or factors were analyzed, one- or two-way ANOVA followed by a Holm-Sidak test for normally distributed groups or a Mann-Whitney test was used for the not normally distributed. Permanova test was used to infer statistically significant differences for the microbiota analysis. The presence of Differentially Abundant (DA) species was assessed using the ANCOM-BC tool (v.2.4.0) (113), enabling Holm-Bonferroni p-values adjusted and a maximum number of 100 iterations. The level of significance used was P < 0.05.

### Study approval

All procedures adhered to the Italian Ministry of Health guidelines for the care and maintenance of laboratory animals, and strictly complied with the European Community Directive n. 2010/63/EU on the protection of animals used for scientific purposes. The Italian Ministry of Health approved animal experimentation at the Institute of Pediatric Research (IRP) in Padua, protocol number: 43F3E.12.EXT.6 (104:2019) and 1038_2024-PR (5168F.26).

### Data availability

The datasets generated and/or analyzed during the current study used for data processing and statistical analyses, are available from the corresponding author upon reasonable request.

## Author contributions

G.C. performed behavioral testing, surgeries, immunohistochemistry, and optical microscopy, analyzed the data, prepared the figures, and wrote the manuscript. C.C. and M.T. performed immunohistochemistry and acquired confocal microscopy data for CXCR4. L.V. supervised behavioral testing and surgeries. R.L. contributed to the analysis of muscle samples. D.P. performed microbiota analysis. G.V. performed hematoxylin and eosin staining of muscle samples and sarcomere length evaluation. R.d.G. supervised hematoxylin and eosin staining and sarcomere length evaluation. M.B. supervised microbiota analysis. M.R. supervised immunohistochemistry and confocal microscopy acquisition for CXCR4. M.A. and G.D. supervised behavioral testing, surgeries, immunohistochemistry, and optical microscopy, analyzed the data, prepared the figures, and wrote the manuscript.

All authors reviewed and approved the final version of the manuscript.

