## Supplementary figures for "Perinatal ischemic stroke impairs motor and cognitive development, muscle structure, and gut microbiota composition, with functional improvement following fecal microbiota transplantation in a mouse model"

**a**

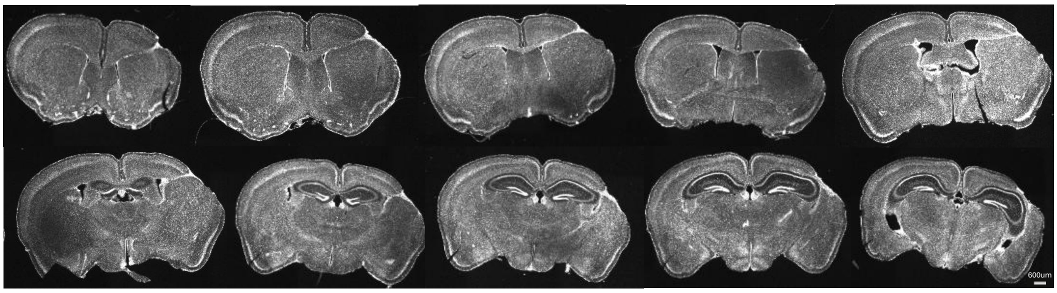


**b**

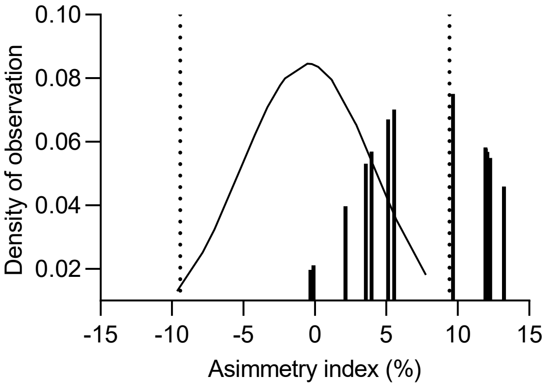


**c**

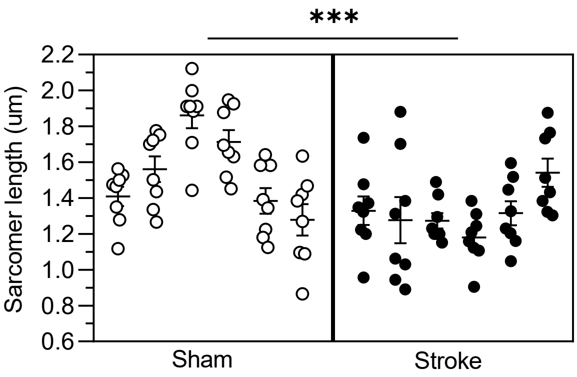


Figure 9

**Figure 9. Supplementary**

(**a**) Representative image of the coronal sections of the brain mouse slice, 79 days after a perinatal stroke with the lesion. Scale bar, 200 μm. (**b**) Distribution of the asymmetry index of the Stroke animals with good and poor recoverers at D79, falling within or outside the threshold, respectively. The dotted line indicates the threshold set at two standard deviations from the mean of the Sham mice. (**c**) Distribution of the different measurements of sarcomere length per animal at D79
